# LuxHMM: DNA methylation analysis with genome segmentation via Hidden Markov Model

**DOI:** 10.1101/2022.12.20.521327

**Authors:** Maia H. Malonzo, Harri Lähdesmäki

## Abstract

DNA methylation plays an important role in studying the epigenetics of various biological processes including many diseases. Although differential methylation of individual cytosines can be informative, given that methylation of neighboring CpGs are typically correlated, analysis of differentially methylated regions is often of more interest.

We have developed a probabilistic method and software, LuxHMM, that uses hidden Markov model (HMM) to segment the genome into regions and a Bayesian regression model, which allows handling of multiple covariates, to infer differential methylation of regions. Moreover, our model includes experimental parameters that describe the underlying biochemistry in bisulfite sequencing and model inference is done using either variational inference for efficient genome-scale analysis or Hamiltonian Monte Carlo (HMC).

Analyses of real and simulated bisulfite sequencing data demonstrate the competitive performance of LuxHMM compared with other published differential methylation analysis methods.

## Background

DNA methylation is an important epigenetic modification associated with many biological processes including various diseases. In promoters, DNA methylation tends to repress gene expression whereas in intragenic locations they tend to upregulate expression [6]. Bisulfite sequencing, whether whole genome (WGBS) or reduced representation (RRBS) bisulfite sequencing, allows for interrogation of DNA methylation at the level of individual CpGs. Moreover, decreasing costs of sequencing have increased the use of these methods. DNA methylation are often studied by analyzing differentially methylated loci (DML) or regions (DMR). Although single differentially methylated CpGs are informative, often DMRs are of more interest [14]. Further, analyzing the combined methylation differences of CpGs within regions increase the statistical power of differential methylation detection.

Given such interest in DMRs, several methods have been developed for identifying them (Table 1). RADMeth uses the beta-binomial regression method in handling complex experimental designs [4]. Beta-binomial regression is used to individually fit single cytosines and then measures the significance of differential methylation using the log-likelihood ratio test between the full and reduced models which generates p-values. To combine information from neighboring cytosines into regions it transforms p-values using the weighted Z-test which then determines which cytosines are combined into regions using an FDR threshold. A method called metilene first recursively segments the genome into regions using the circular binary segmentation algorithm which generates regions that maximizes the difference of CpG-wise mean methylation levels [7]. Then, it calculates p-values using a version of the Kolmogorov-Smirnov test which tests the significance of potential DMRs. HMM-DM uses hidden Markov model (HMM) to segment the genome into regions and Bayesian methods to infer model parameters. It then uses MCMC to compute the posterior probability of each state: hypermethylated, equally methylated or hypomethylated. To identify DMRs, it joins hypermethylated or hypomethylated CpGs into regions. In DMRcate, standard linear modelling is performed using limma which generates a signed statistic for measuring the difference between treatment effects per CpG site [13]. The square of this value is then applied to a Gaussian smoother. It then uses an approximation that generates a value for which a p-value is computed by comparison to a chi-square distribution. Individual sites below a given p-value threshold are selected and grouped into regions that are separated by, at most, a threshold number of nucleotides. DSS models the methylation counts by a beta-binomial distribution with an arcsine link function and fits the transformed methylation levels with a generalized least squares procedure from which it obtains estimates of the model coefficients at each CpG site [12]. Hypothesis testing is performed using Wald test on the coefficient estimates. After detection of statistically significant CpG sites, DSS merges nearby loci into regions.

**Table 1.** Methods comparison.

| Method | Methylation model | Algorithm for CpG correlation |
| --- | --- | --- |
| RADMeth | beta-binomial | weighted Z-test |
| metilene | Kolmogorov-Smirnov | circular binary segmentation |
| HMM-DM | Bayesian HMM | HMM |
| DMRcate | linear Gaussian (limma) | kernel smoothing |
| DSS | beta-binomial | smoothing via moving average |
| LuxUS | Bayesian GLMM | random effect correlation structure |
| LuxHMM | Bayesian GLM | HMM |

LuxGLM [1] and LuxUS [5] use extended versions of generalized linear model (GLM) to analyze methylation data with complex experimental designs and incorporate estimation of experimental parameters that describe the underlying biochemistry in methylation sequencing data. LuxGLM uses matrix normal distribution to handle multiple methylation modifications. LuxUS uses a generalized linear mixed model (GLMM) to analyze cytosines within a genomic window simultaneously. To analyze the spatial correlation of cytosines it uses a random effect correlation structure. It also analyzes the variation of individual replicates using a replicate random effect. Features of previous methods as well as the proposed method, LuxHMM, are contrasted in Table 1.

## Implementation

Bisulfite sequencing data consists of DNA where unmethylated cytosines are converted into uracil by bisulfite treatment and sequenced as thymine to differentiate it from methylated cytosine which are not converted and sequenced as cytosine.

A commonly used methylation level estimate is obtained by taking the ratio of methylated cytosine to the sum of methylated and unmethylated cytosine, *µ* = *N*_BS,C_*/N*_BS_. To infer differential methylation, the methylation levels between groups are compared. Hypermethylation occurs when the methylation level for a comparison (or treatment) group is generally higher compared to a reference (or control) group, and hypomethylation when it is lower. We are interested in modeling methylation levels and differential methylation across *T* cytosines *c*_1_, *c*_2_, …, *c*_*T*_. Differentially methylated regions are often of more interest than single cytosines due to their combined effect compared to the individual effect of a single cytosine. A methylated region *C* consists of consecutive CpGs *c*_*t*_s that are hypermethylated, hypomethylated or have equal methylation (*M*_*j*_), *C* = {*c*_*t*_ |*c*_*t*_ ∈ *M*_*j*_}. A region is differentially methylated when it is either hypermethylated or hypomethylated.

Our method consists of two modules: 1) genome segmentation via HMM, and 2) estimation of methylation levels and inference of differential methylation using Bayesian GLM. In inference of differential methylation, significance of explanatory variable is measured by Bayes factors.

### Genome segmentation via HMM

To extract regions from a sequence of cytosines, we use hidden Markov model (HMM). HMM is a statistical model that infers a sequence of hidden states from a sequence of observations. In this work, the hidden states *x* are the methylation states, specifically: 1) hypermethylation, 2) hypomethylation, and 3) equal methylation between two groups. For each cytosine, the observations *y* are the differences in the mean methylation levels between groups, 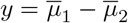, where 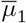 is the mean methylation level for one group and 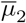 for another.

HMM is parameterized by two distributions: the observation emission probabilities and the state transition probabilities (Fig. 1). The observation emission probabilities, *P* (*y*_*t*_|*x*_*t*_), give the probability of observing *y* at cytosine position *t* given the underlying hidden state *x*_*t*_, i.e. the probability of observing the differences in methylation levels between two groups given the underlying methylation states *M*_*j*_ (hypermethylation, hypomethylation or equal methylation). The state transition probabilities, *P* (*x*_*t*_|*x*_*t−*1_), give the probability of hidden state *x*_*t−*1_ moving to *x*_*t*_ in a sequence, i.e. the probability of moving from one methylation state to another (or remaining the same) between two consecutive CpGs.

**Figure 1.**
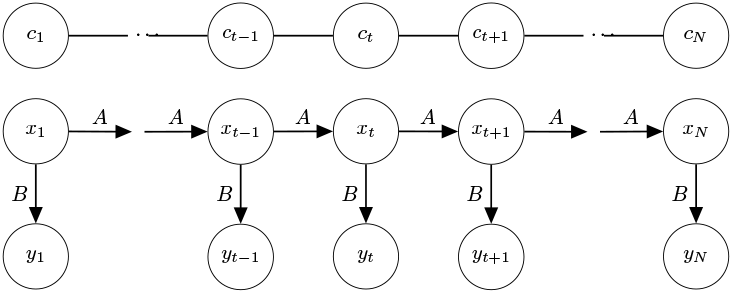
Diagram of emission and transition probabilities. The top-most row (*C*) indicates the cytosine position, the second row (*X*) denotes the hidden methylation states and the bottom row (*Y*) represents the observed differences in methylation levels between groups. *A* denotes the state transition probabilities and *B* the observation emission probabilities.

For a given hidden state sequence *X* = *x*_1_, *x*_2_, …, *x*_*T*_ and observation sequence *Y* = *y*_1_, *y*_2_, …, *y*_*T*_, the observation sequence likelihood is

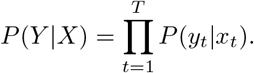

It is straightforward to compute the joint probability of a given sequence of methylation states and a sequence of observed methylation differences

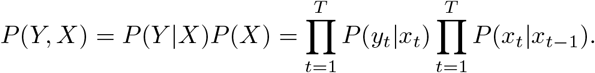

The total probability of the observed methylation differences can then be obtained by summing over the hidden states

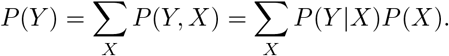

With these definitions we can select the hidden state sequence that maximizes the observation likelihood. However, this is infeasible due to the high number of possible state sequences. Instead a dynamic programming algorithm like the Viterbi algorithm recursively computes *v*_*t*_(*j*) which denotes the probability of being in state *j* given the observations for the first *t* cytosines. For a given state *x*_*j*_ at cytosine position *t, v*_*t*_(*j*) is computed by

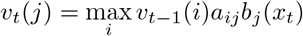

where *v*_*t−*1_(*i*) is the previous Viterbi path probability from the previous time step, *a*_*ij*_ is the transition probability from previous state *x*_*i*_ to current state *x*_*j*_ and *b*_*j*_(*y*_*t*_) is the emission probability of the observation *y*_*t*_ given state *j* [8].

To learn the most likely transition, ***A*** *= {a*_*ij*_}, and emission, **B** = {*b*_*j*_(*y*_*t*_)}, probabilities and initial state distribution *π*_*i*_ = *P* (*X*_1_ = *i*), the Baum-Welch algorithm, another dynamic programming algorithm, finds a (local) maximum of *θ* = arg max_*θ*_ *P* (*Y* | *θ*), where *θ* = (*A, B, π*), using the expectation-maximization (EM) algorithm [2].

In this work we use pomegranate, a Python package that implements probabilistic models, including HMMs [15]. The model is initialized with state and transition probabilities. We assume the emission distributions follow a Gaussian distribution *𝒩*(*ψ, σ*^2^), where *ψ* and *σ*^2^ are set to 0 and 0.08 (equal methylation), 0.3 and 0.06 (hypermethylation) and -0.3 and 0.06 (hypomethylation). The transition probabilities were optimized using the Baum-Welch algorithm using the initial values shown in Supplementary Section 1.

To determine the most likely sequence of hidden states, i.e. the sequence of methylation states, we use the Viterbi algorithm implemented in the package. To learn the most likely emission and transition probabilities given the sequence of observations we use the Baum-Welch algorithm, also supported by pomegranate. After learning the hidden methylation states, adjacent cytosines with the same methylation state are combined into regions, as well as the total read counts 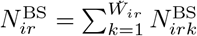, where *k* is the *k*th CpG in *C*_*ir*_ and *W*_*ir*_ = |*C*_*ir*_| is the number of consecutive CpGs with the same methylation state in the *i*th sample and the *r*th region and, similarly for methylated read counts,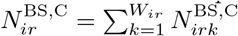.

### Estimation of methylation levels and differential methylation

We briefly review the underlying statistical model for the experimental parameters [1]. Experimental parameters that define the underlying biochemistry in bisulfite sequencing should be considered in estimation of methylation levels. Bisulfite conversion rate (BS_eff_), sequencing error (seq_err_) and incorrect bisulfite conversion rate 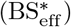 can significantly affect methylation estimates. Low BS_eff_ causes overestimation of methylation levels whereas high 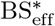 results in underestimation. On the other hand, high seq_err_ can lead to either overestimation or underestimation. BS_eff_ can be estimated by using the lambda phage genome. Since the lambda phage genome is unmethylated, BS_eff_ can be estimated by taking the ratio of all cytosine reads converted into thymine over the total number of reads. Similarly, 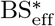 can be estimated with spike-ins of oligonucleotides where all the cytosines are methylated. On the other hand, seq_err_ can be estimated using Phred scores *Q* by converting them to base-calling error probabilities 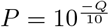.

Given the above definitions, 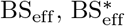 and seq_err_ determine the conditional probability of a sequencing readout being “C”, given that the cytosine is methylated or unmethylated (Fig. 2). Specifically, since BS_eff_ is the probability of an unmethylated cytosine being converted into uracil, 1− BS_eff_ is the probability of an unmethylated cytosine incorrectly not converted into uracil. If an unmethylated cytosine is correctly converted into uracil it still has seq_err_ probability of being incorrectly sequenced as “C”. Whereas, if it is incorrectly not converted to uracil and remains a cytosine, it has 1− seq_err_ probability of being correctly sequenced as “C”. Put together, the conditional probability of sequencing “C” given the cytosine is unmethylated is

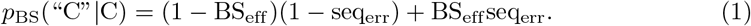

**Figure 2.**
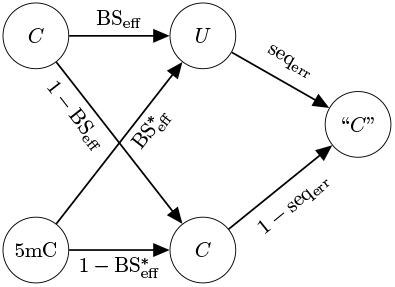
Probability tree of observing “C” readout when the true methylation state is methylated or unmethylated.

On the other hand, if a cytosine is methylated, the probability that it is correctly not converted to uracil is 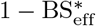 and the probability that it is correctly sequenced as “C” is 1 − seq_err_. The probability that the unmethylated cytosine is incorrectly converted to uracil and incorrectly sequenced as “C” are, respectively, 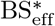 and seq_err_. Thus, the conditional probability of sequencing “C” given the cytosine is methylated is

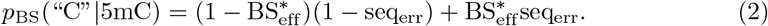

Thus far we have described individual cytosines. However, this description can be generalized to DNA regions. Let *θ* ∈ [0, 1] represent the unknown fraction (or probability) of methylated DNA. Following Eqs. 1 and 2, the probability of observing “C” readouts for a given region is *p*_BS_(“C”) = *p*_BS_(“C” | 5mC)*θ* + *p*_BS_(“C” |C)(1 −*θ*). Finally, the total number of “C” readouts is binomially distributed,

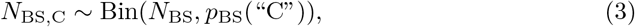

where *N*_BS_ is the total number of reads. See Fig. 3 for the plate diagram of the model.

**Figure 3.**
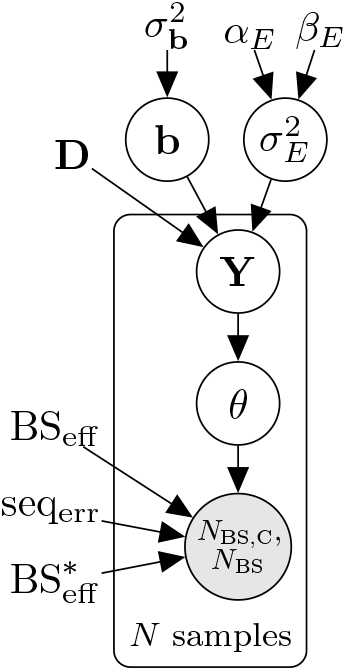
Plate diagram of the LuxHMM model for analyzing experimental parameters and methylation levels. The circles represent latent (white) and observed (gray) variables and the unbordered nodes represent hyperparameters and constant values.

To incorporate complex experimental designs to the model, we simplify the method proposed in [5] by doing away with the spatial correlation component and use generalized linear regression,

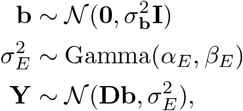

where 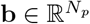 (where *N*_*p*_ is the number of covariates, possibly including the intercept) is the vector of regression coefficients, 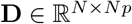 is the design matrix, and **Y** ∈ R*N*. The values of the hyperparameters are 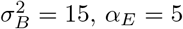, and *β*_*E*_ = 5, and were taken from [5]. We apply this model to regions instead of single CpGs to speed up computation. Finally, we use the sigmoid link function

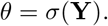

The model is implemented using the probabilistic programming language Stan [3], and model inference is done using either Hamiltonian Monte Carlo (HMC) or automatic differentiation variational inference (ADVI) for faster estimation of the model parameters [10], both built-in features of Stan. Stan uses a locally adaptive version of dynamic HMC sampling. In variational inference (VI) the posterior *p*(*ϕ*| *D*) of all unknowns *ϕ* given observed data *D* is approximated with a simpler distribution *q*(*ϕ*; *ρ*), which is selected from a chosen family of distributions by minimizing divergence between *p*(*ϕ*|*D*) and *q*(*ϕ*; *ρ*).

To detect differential methylation w.r.t. any of the *N*_*p*_ covariates in **D**, hypothesis testing was done using Bayes factors via the Savage-Dickey density ratio method as implemented in [1].

## Results

To demonstrate the performance of LuxHMM and how well it performs compared to other methods, we analyze real and simulated BS-seq datasets. The first dataset is a simulated dataset based on real BS-seq data. The second is a simulated BS-seq dataset generated using a general experimental design. Lastly, we use a real BS-seq dataset with confounding covariates. We compare the performance of LuxHMM with RADMeth, metilene, HMM-DM, DMRcate and DSS.

### Comparison of performance on simulated dataset based on real BS-seq data

To assess the accuracy of our method compared to other published methods we used a simulated dataset by [9]. Bisulfite sequencing data was obtained from real CpG islands which allowed variance and correlation to be incorporated into the simulated dataset. The dataset was derived from 12 individuals which were divided into 6 controls and 6 cases. The dataset was divided into two sets wherein 10,000 DMRs were incorporated into one set. Methylation counts were added to or substracted from the case samples so that the methylation differences were 0.1, 0.2, 0.3 or 0.4.

In LuxHMM, either all regions or only hypo- and hypermethylated regions, as classified by HMM, were used as input in determining DMRs. Parameter settings for competing methods are described in Supplementary Section 2.

The area under the receiver operating curve (AUROC) and the average precision (AP), to handle the imbalance in the dataset given that there are much more negative than positive samples, were computed (Table 2). For AP, the baseline is 0.11 which is the fraction of the number of true positives over the total number tests. True positives are differentially methylated cytosines whereas negatives are non-differentially methylated cytosines. In all methods, cytosines which are not covered by the returned regions are given a score of zero. The highest AUROC and AP were generated by LuxHMM used with all regions. The higher recall suggests that the state assignment of HMM misses differentially methylated regions which are inaccurately classified as regions with equal methylation between two groups. This also demonstrates that LuxHMM more accurately detects DMRs compared to the other methods used. Another notable result is that DMRcate has a relatively high AUROC and a low AP. This could be caused by a high false positive rate which is masked in AUROC due to a high number of true negative samples. As true negative samples are excluded in the computation of AP, the high false positive rate results in a low AP.

**Table 2.** AUROC and AP for simulated dataset from [9].

| Method | AUROC | AP |
| --- | --- | --- |
| LuxHMM <sup>1</sup> | **0.946** | **0.851** |
| LuxHMM <sup>2</sup> | 0.933 | 0.834 |
| LuxUS | 0.889 | 0.606 |
| RADMeth | 0.847 | 0.682 |
| metilene | 0.833 | 0.686 |
| HMM-DM | 0.633 | 0.357 |
| DMRcate | 0.832 | 0.271 |
| DSS | 0.855 | 0.723 |

<sup>1</sup> All regions
<sup>2</sup> Hypo- and hypermethylated regions

### Alternative emission and initial transition probabilities

To test the sensitivity of the proposed model to different emission distribution parameters, we tested various parameter values on the [9] dataset using all regions. Table 3 shows that the model is not sensitive to different values of standard deviation but is sensitive to the means, with the highest AP when using means -0.3 and 0.3.

**Table 3.** AUROC and AP for different emission distributions.

| Equal |  | Hypo |  | Hyper |  | AUROC | AP |
| --- | --- | --- | --- | --- | --- | --- | --- |
| mean | std. dev. | mean | std. dev. | mean | std. dev. |  |  |
| 0 | 0.08 | -0.1 | 0.06 | 0.1 | 0.06 | 0.932 | 0.735 |
| 0 | 0.06 | -0.3 | 0.04 | 0.3 | 0.04 | 0.946 | 0.852 |
| 0 | 0.06 | -0.3 | 0.05 | 0.3 | 0.05 | 0.946 | 0.852 |
| 0 | 0.08 | -0.3 | 0.06 | 0.3 | 0.06 | 0.946 | 0.851 |
| 0 | 0.1 | -0.3 | 0.07 | 0.3 | 0.07 | 0.946 | 0.852 |
| 0 | 0.1 | -0.3 | 0.08 | 0.3 | 0.08 | 0.946 | 0.852 |
| 0 | 0.1 | -0.3 | 0.1 | 0.3 | 0.1 | 0.946 | 0.852 |
| 0 | 0.08 | -0.5 | 0.06 | 0.5 | 0.06 | 0.918 | 0.795 |
| 0 | 0.1 | -0.5 | 0.07 | 0.5 | 0.07 | 0.917 | 0.796 |

We also tested using five hidden states with two hidden states each for the hypo- and hypermethylated regions (Table 4). The AUROC and AP are, respectively, 0.948 and 0.846, indicating that increasing the number of hidden states from three to five does not increase accuracy.

**Table 4.**
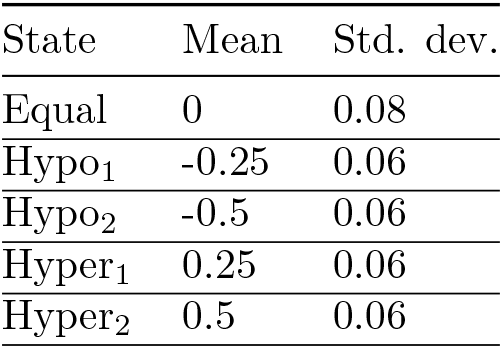
Emission parameters for a HMM model with five hidden states.

When not specifying the emission distributions and letting pomegranate instead estimate the emission distributions we obtain a higher AUROC and a lower AP (Table 5). We prioritize AP as it takes into account the imbalanced dataset. Genome segmentation was based on Fig. 4. For three hidden states, we used *s*1 as hypo- and hypermethylated states (with *s*0 and *s*2 as states with no difference between groups), whereas with four hidden states we used *s*1 and *s*2 as hypo- and hypermethylated states (with *s*0 and *s*3 as states with no difference between groups). In computing AUROC and AP we used either all hidden states (including state with no difference between groups) or just hypo- and hypermethylated states.

**Table 5.** AUROC and AP when not specifying state distributions.

| Number of states | AUROC | AP |
| --- | --- | --- |
| $3^1$ | 0.938 | 0.753 |
| $4^1$ | 0.934 | 0.768 |
| $3^2$ | 0.965 | 0.790 |
| $4^2$ | 0.973 | 0.815 |

1 All regions
2 Hypo- and hypermethylated regions

**Figure 4.**
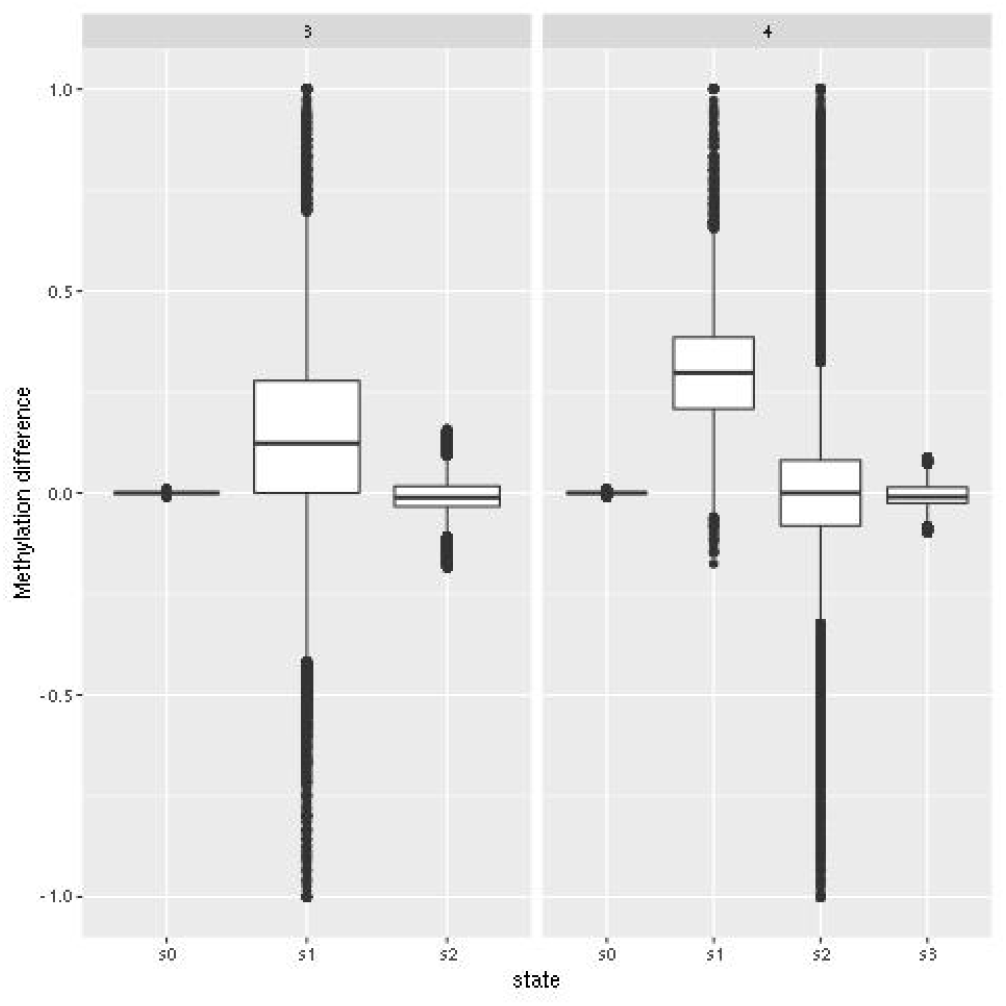
Distribution of methylation differences with three and four hidden states when the emission distributions are not specified.

### Comparison of performance on simulated dataset with confounding covariates

To test the performance of LuxHMM in datasets with general experimental design we simulated a dataset with multiple covariates: 1) binary case/control, 2) arbitrary binary, 3) arbitrary continuous. The design matrix **D** is shown in Table 6. This simulation was modified from [13].

**Table 6.** Design matrix for simulated data.

| Intercept | Case/control | Binary | Continuous |
| --- | --- | --- | --- |
| 1 | 0 | 0 | 0.3 |
| 1 | 0 | 1 | 0.5 |
| 1 | 0 | 0 | 0.7 |
| 1 | 1 | 1 | 0.3 |
| 1 | 1 | 0 | 0.5 |
| 1 | 1 | 1 | 0.7 |

To model the varying lengths of methylated regions, the length *L* of the regions in terms of number of CpGs was sampled from *L*∼ ceiling(gamma(shape = 4, rate = 0.2)). The genomic coordinates were taken from the hg19 build. To model the varying differences in methylation levels, the covariate coefficients **b** were sampled from **b**∼ *𝒩* (*µ* = 0, *σ*^2^ = 5). For non-differentially methylated regions, the coefficient corresponding to the covariate of interest was set to zero. Conversely, for differentially methylated regions, the coefficient corresponding to the covariate of interest *b* was set so that *b <* −3 or *b >* 3 to ensure significant differential methylation. Finally, *θ* = *σ*(**Y**) where 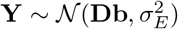 where 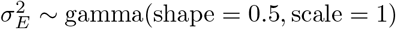 (shape = 0.5, scale = 1). Around 1,700 DMRs were added to the genome.

In LuxHMM, either all regions or only hypo- and hypermethylated regions, as classified by HMM, were used as input in determining DMRs. Parameter settings for competing methods are described in Supplementary Section 3.

AUROC and AP, to handle the imbalance in the dataset given that there are much more negative than positive samples, were computed (Table 7). For AP, the baseline is 0.0014. LuxHMM using all regions generated the highest AUROC and LuxHMM using just hypo- and hypermethylated regions generated the highest AP. This indicates that, like in Section, using LuxHMM with all regions has a higher recall whereas using LuxHMM with just hypo- and hypermethylated regions has a higher precision. This also shows that LuxHMM is able to more accurately detect DMRs from a dataset with confounding covariates.

**Table 7.** AUROC and AP for simulated dataset with confounding covariates.

| Method | AUROC | AP |
| --- | --- | --- |
| LuxHMM <sup>1</sup> | **0.823** | 0.536 |
| LuxHMM <sup>2</sup> | 0.756 | **0.549** |
| LuxUS | 0.679 | 0.321 |
| RADMeth | 0.644 | 0.246 |
| metilene | 0.714 | 0.348 |
| HMM-DM | 0.616 | 0.180 |
| DMRcate | 0.658 | 0.065 |
| DSS | 0.672 | 0.339 |

<sup>1</sup> All regions<sup>2</sup> Hypo- and hypermethylated regions

### Comparison of performance on real BS-seq data with confounding covariates

To test the performance of LuxHMM on real BS-seq data with multiple covariates we evaluated the different statistical methods in terms of gene set enrichment using the webtool GREAT [11] on the dataset with GEO accession number GSE47966 as originally performed by [16]. The dataset consists of samples taken from mice brain tissue (WGBS). Three samples consisted of neuron cells and three consisted of non-neuron cells. In addition, the samples were divided into male and female mice and different ages (6 week and 12 month old females, and 7 week old males). DMRs between neurons and non-neurons were identified using the different methods and then gene ontology (GO) enrichment were performed to test the ability of the various methods to identify biologically relevant regions. The top 25 and 60 enriched GO terms based on binomial ranking were taken and the percentage of GO terms related to the neural system were determined. Gene set enrichment analysis were performed with mouse phenotype annotations.

In LuxHMM, hypo- and hypermethylated regions, as determined by HMM, were used as input in determining differentially methylated regions. HMC was used to sample from the posterior distribution with four chains, 1,000 iterations for warmup for each chain and a total of 1,000 iterations for sampling. In addition, as in [16], for the regions, a threshold of *>* 25 CpGs was used. To make a comparable assessment, the top 10,000 to 15,000 DMRs from all methods were used as input to GREAT. Parameter settings for competing methods are described in Supplementary Section 4.

As shown in Table 8, HMM-DM generated the highest percentages of enriched GO terms related to the neural system in both the top 25 and top 60 enriched GO terms. In the top 25 enriched GO terms, LuxHMM generated the second highest number of enriched GO terms related to the neural system and in the top 60 LuxHMM was fourth highest after DSS and LuxUS (Supplementary file). This shows that LuxHMM performs comparatively well in finding biologically relevant regions relative to other methods tested.

**Table 8.** Enriched GO terms related to the neural system.

| Method | Top 25 (%) | Top 60 (%) |
| --- | --- | --- |
| LuxHMM <sup>1</sup> | 92 | 83 |
| LuxUS | 88 | 85 |
| RADMeth | 88 | 80 |
| metilene | 20 | 32 |
| HMM-DM | **96** | **93** |
| DMRcate | 84 | 68 |
| DSS | 88 | 87 |

<sup>1</sup> Hypo- and hypermethylated regions

## Conclusions

We propose the tool LuxHMM for detecting differentially methylated regions. This tool uses HMM to segment the genome into regions with hypomethylation, hypermethylation and equal methylation between two groups and Bayesian regression for evaluating differential methylation. Further, model inference is done using either variational inference for efficient genome-scale analysis or HMC.

We show using simulated and real BS-seq data with general experimental designs that LuxHMM outperforms other published methods in detecting differentially methylated regions from simulated datasets and performs comparatively well in a real dataset.

## Acknowledgments

We acknowledge the computational resources provided by the Aalto Science-IT project.

## Author’s contributions

MM and HL developed the method. MM implemented the method. MM and HL wrote the manuscript. All authors read and approved the final version of the manuscript.

## References

1. T. Äijö, X. Yue, A. Rao, and H. Lähdesmäki. Luxglm: a probabilistic covariate model for quantification of dna methylation modifications with complex experimental designs. Bioinformatics, 32:i511–i519, 2016.

2. J. A. Bilmes. A Gentle Tutorial of the EM Algorithm and its Application to Parameter Estimation for Gaussian Mixture and Hidden Markov Models. Berkeley, CA: International Computer Science Institute, Berkely, CA, 1998.

3. B. Carpenter, A. Gelman, M. D. Hoffman, D. Lee, B. Goodrich, M. Betancourt, M. Brubaker, J. Guo, P. Li, and A. Riddell. Stan: A probabilistic programming language. Journal of Statistical Software, 76, 2017.

4. E. Dolzhenko and A. D. Smith. Using beta-binomial regression for high-precision differential methylation analysis in multifactor whole-genome bisulfite sequencing experiments. BMC bioinformatics, 15:1–8, 2014.

5. V. Halla-Aho and H. Lähdesmäki. Luxus: Dna methylation analysis using generalized linear mixed model with spatial correlation. Bioinformatics, 36:4535–4543, 2020.

6. J. Jeschke, C. E, and F. F. Dna methylome profiling beyond promoters-taking an epigenetic snapshot of the breast tumor microenvironment. The FEBS journal, 282:1801–1814, 2015.

7. F. Juhling, H. Kretzmer, S. H. Bernhart, C. Otto, P. F. Stadler, and S. Hoffmann. Metilene: fast and sensitive calling of differentially methylated regions from bisulfite sequencing data. Genome research, 26:256–262, 2016.

8. D. Jurafsky and M. J. H. Speech and language processing, 2021.

9. H. U. Klein and K. Hebestreit. An evaluation of methods to test predefined genomic regions for differential methylation in bisulfite sequencing data. Briefings in bioinformatics, 17:796–807, 2016.

10. A. Kucukelbir, R. Ranganath, A. Gelman, and D. Blei. Automatic variational inference in stan. In C. Cortes, D. D. Lee, M. Sugiyama, and G. R, editors, Advances in Neural Information Processing Systems 28 (NIPS 2015), pages 568–576. Neural Information Processing Systems, 2015.

11. C. Y. McLean, D. Bristor, M. Hiller, S. L. Clarke, B. T. Schaar, C. B. Lowe, M. Wenger, and G. Bejerano. Great improves functional interpretation of cis-regulatory regions. Nature biotechnology, 28:495–501, 2010.

12. Y. Park and H. Wu. Differential methylation analysis for bs-seq data under general experimental design. Bioinformatics, 32:1446–1453, 2016.

13. T. J. Peters, M. J. Buckley, A. L. Statham, R. Pidsley, K. Samaras, R. V. Lord, S. J. Clark, and P. L. Molloy. De novo identification of differentially methylated regions in the human genome. Epigenetics & chromatin, 8:1–16, 2015.

14. M. D. Robinson, A. Kahraman, C. W. Law, H. Lindsay, M. Nowicka, L. M. Weber, and X. Zhou. Statistical methods for detecting differentially methylated loci and regions. Frontiers in genetics, 5:324, 2014.

15. J. Schreiber. Pomegranate: fast and flexible probabilistic modeling in python. Journal of Machine Learning Researc, 18:1–6, 2018.

16. Y. Wen, F. Chen, Q. Zhang, Y. Zhuang, and Z. Li. Detection of differentially methylated regions in whole genome bisulfite sequencing data using local getis-ord statistics. Bioinformatics, 32:3396–3404, 2016.

