## Supplemental material for "LuxHMM: DNA methylation analysis with genome segmentation via Hidden Markov Model"

### Supplementary information

Maia Malonzo & Harri Lähdesmäki

December 21, 2022

#### 1 Initial state transition probabilities

Table 1: Initial state transition probabilities

| | Hyper | Hypo | Equal |
|---|---|---|---|
| Hypermethylation | 0.50 | 0.25 | 0.25 |
| Hypomethylation | 0.25 | 0.50 | 0.25 |
| Equal methylation | 0.25 | 0.25 | 0.50 |

#### 2 Parameter settings for competing methods on simulated dataset based on real BS-seq data

For RADMeth we used bins 1:100:1 and 1:200:50 in combining significance of CpGs. This parameter determines the distance at which correlation of p-values is computed. Using bins 1:100:1 returned a higher AUROC and AP hence we present the result with this setting. We used a p-value cutoff of 0.05 which joins neighboring differentially methylated CpGs with p-values below this cutoff. For computing AUROC and AP we use the log odds ratio as the metric for scoring regions. In metilene, we use the default parameters including the maximum adjusted p-value at 0.05. In computing AUROC and AP, $1 - q$-value is used as the metric for scoring regions. In HMM-DM, default parameters were used except min.percent which was set to 0.4 as this maximizes AUROC and AP. The mean posterior probability was used as metric for scoring in AUROC and AP. In DMRcate, default settings were used, including an FDR threshold at 0.05 in filtering CpGs. The metric used to score the returned regions was $1 - \mathrm{HMFDR}$ (harmonic mean of the individual component FDRs). For DSS, default settings were used and p.threshold of 0.05. The metric used to score the regions was areaStat defined as sum of the test statistics of all CpG sites within the DMR.

DSS was run with and without smoothing (with spans equal to 500bp and 1000bp). Without smoothing generated higher AURCOC and AP so these were included in the comparison. LuxUS was run with ADVI and the metric used to score the DMRs was Bayes factors. The number of output samples, the evidence lower bound (ELBO) samples and grad samples were 1000, 1000 and 10, respectively. These values were previously found to perform well in Malonzo *et al.* (2022).

### 3 Parameter settings for competing methods on simulated dataset with confounding covariates

For comparing LuxHMM with other methods, we use the same pipeline used in Section 3.1. For RADMeth we used bins 1:100:1 and 1:200:50 in combining significance of CpGs. Using bins 1:200:50 returned a higher AUROC and AP hence we present the result with this setting. RADMeth does not handle continuous covariates hence we binarized the continuous-valued covariate. We used a p-value cutoff of 0.05. In metilene, we use the default parameters including the maximum adjusted p-value at 0.05. metilene does not handle multiple covariates so the dataset is only divided between controls and cases. In HMM-DM, default parameters were used except min.percent which was set to 0.4 as this was previously shown to maximize AUROC and AP in Section 3.1. HMM-DM does not handle multiple covariates so the dataset was only divided into case and control samples. In DMRcate, default settings were used. The FDR threshold at which individual CpGs are called was set to 0.10 and 0.05. The latter generated a higher AP hence we present the results using the FDR threshold at 0.05. For DSS default settings were used and p.threshold of 0.05. DSS was run with and without smoothing (with span equal to 500bp). Without smoothing generated a higher AP so this was included in the comparison. LuxUS was run with ADVI.

### 4 Parameter settings for competing methods on real BS-seq data with confounding covariates

For comparing LuxHMM with other methods, we use the same pipeline used in Section 3.1. For RADMeth we used bins 1:100:1 in combining significance of CpGs. RADMeth does not handle continuous covariates hence we binarized the age covariate. In metilene, we use the default parameters. metilene does not handle multiple covariates so the dataset is only divided between neurons and non-neurons. In HMM-DM, default parameters were used except min.percent which was set to 0.4 as this was previously shown to maximize AUROC and AP in Section 3.1. HMM-DM does not handle multiple covariates so the dataset was only divided into neurons and non-neurons. In DMRcate, default settings were used. For DSS default settings were also used. DSS was run without smoothing as this was previously found to perform better in Section 3.1. LuxUS was run with HMC for evaluating the posterior distribution. The number of chains was four and the total number of posterior samples was 1,000. The burn-in period consisted of 1,000 samples for each chain.

### References


Malonzo, M. H., Halla-Aho, V., Konki, M., Lund, R. J., & Lähdesmäki, H. (2022). LuxRep: a technical replicate-aware method for bisulfite sequencing data analysis. *BMC bioinformatics*, **23**(1):1-19.
